# Directed networks as a novel way to describe and analyze cardiac excitation: Directed Graph mapping

**DOI:** 10.1101/596288

**Authors:** Nele Vandersickel, Enid Van Nieuwenhuyse, Nico Van Cleemput, Jan Goedgebeur, Milad El Haddad, Jan De Neve, Anthony Demolder, Teresa Strisciuglio, Mattias Duytschaever, Alexander V. Panfilov

## Abstract

Networks provide a powerful methodology with applications in a variety of biological, technological and social systems such as analysis of brain data, social networks, internet search engine algorithms, etc. To date, directed networks have not yet been applied to characterize the excitation of the human heart. In clinical practice, cardiac excitation is recorded by multiple discrete electrodes. During (normal) sinus rhythm or during cardiac arrhythmias, successive excitation connects neighboring electrodes, resulting in their own unique directed network. This in theory makes it a perfect fit for directed network analysis. In this study, we applied directed networks to the heart in order to describe and characterize cardiac arrhythmias. Proofof-principle was established using in-silico and clinical data. We demonstrated that tools used in network theory analysis allow to determine the mechanism and location of certain cardiac arrhythmias. We show that the robustness of this approach can potentially exceed the existing state-of-the art methodology used in clinics. Furthermore, implementation of these techniques in daily practice can improve accuracy and speed of cardiac arrhythmia analysis. It may also provide novel insights in arrhythmias that are still incompletely understood.

## 1. Introduction

A network, in the most general sense, is a collection of nodes connected by links, which can represent diverse systems. Over the past twenty years network theory has had many applications, ranging from biology to social sciences (1). Examples include the PageRank algorithm (2) for the World Wide Web which formed the basis of Google, determining the shortest route(s) between two places, modelling of molecules (e.g. fullerenes (3)), social networks (4), interactions of genes, proteins, metabolites and other cellular components (5, 6), the spread of diseases (7, 8), and many others. More recently, networks led to the development of novel diagnostic biomarkers in Alzheimer’s disease, multiple sclerosis, traumatic brain injury and epilepsy (9). A network can be directed or undirected, depending if the connecting links have a direction from one node to the other. In spite of this diversity, directed networks have not yet been applied to describe the excitation of the heart. Cardiac contraction is mediated by electrical waves of excitation and abnormal propagation of these waves leads to cardiac arrhythmias. Cardiac arrhythmias remain a major challenge in health care. Among the most common types of cardiac arrhythmias are atrial tachycardia (AT), atrial fibrillation (AF), ventricular tachycardia (VT) and ventricular fibrillation (VF). In recent years, the understanding and treatment of cardiac arrhythmias has improved due to the ability of recording electrical activity from the endo- or epicardial surface of the heart. The most reliable recordings are made by multiple electrodes placed on the myocardium, each giving a local activation time (LAT) of the excitation wave at discrete points of measurements. As such, the cardiac excitation network is suitable for directed network analysis by connecting neighboring points according to their LAT values.

In clinical electrophysiology it is important to determine the driving mechanism of an arrhythmia and its location. There are two main driving mechanisms for cardiac arrhythmias. The first mechanism is reentry, which is characterized by rotational activation. Reentry can be anatomical or functional. Anatomical reentry is characterized by circular electrical propagation around an obstacle (e.g. scar tissue). The size of the smallest rotation allows for further differentiation between macro-reentry and localized reentry. Functional reentry is characterized by reentry around an excitable core (often referred to as rotors (10)). The second driving mechanism of arrhythmias is triggered activity or automaticity. The latter mechanism gives rise to focal activation, with most likely a centrifugal activation pattern.

At present, it remains challenging to precisely determine the mechanism of arrhythmia in a given experimental setup or patient. When the underlying mechanism of the arrhythmia can be determined, the correct ablation target becomes easier to identify, making it possible to cure the patient from the arrhythmia (11). If an incorrect target is ablated, not only will the patient not be cured, but new arrhythmias may be induced (due to scarring). Therefore, it is crucial to identify the correct ablation target. For example, ablation of some ATs can be very complex, because patterns of scar and wavefront propagation are unique in each patient (12). In these cases, the ablation procedure tends to be complex and time-consuming. This is particularly true for ATs occurring after ablation of persistent AF (13). In the present study, directed networks are used to determine rotational and focal activity with the potential of guiding future ablation strategies. We refer to this method as directed graph mapping (DG-mapping). *The goal of this work is to demonstrate the wide applicability of directed networks to the heart for each driving mechanism of cardiac arrhythmia in both the atria and the ventricles*. Therefore, we tested the accuracy of DG-mapping in in-silico (ventricular) models of functional and anatomical reentry and focal activity. To determine the accuracy of DG-mapping in the atria, we analyzed 31 clinical cases of atrial tachycardia. Regular AT is a clinical tachyarrhythmia in which the operator can be sure about the location of the tachycardia since ablation of the correct target almost always results in immediate success. Therefore, AT was used as the gold standard for validating DG-mapping in a clinical setting. In addition, DG-mapping was compared to phase mapping (14) via in-silico simulations, a widely used technique for detecting the center of a rotor.

## 2. Material and Methods

In the next sections the general protocol of DG-mapping is explained according to the flowchart given in Figure 1. Figure 2 and 3 demonstrate DG-mapping on a simulated and a clinical example.

**Fig. 1.**
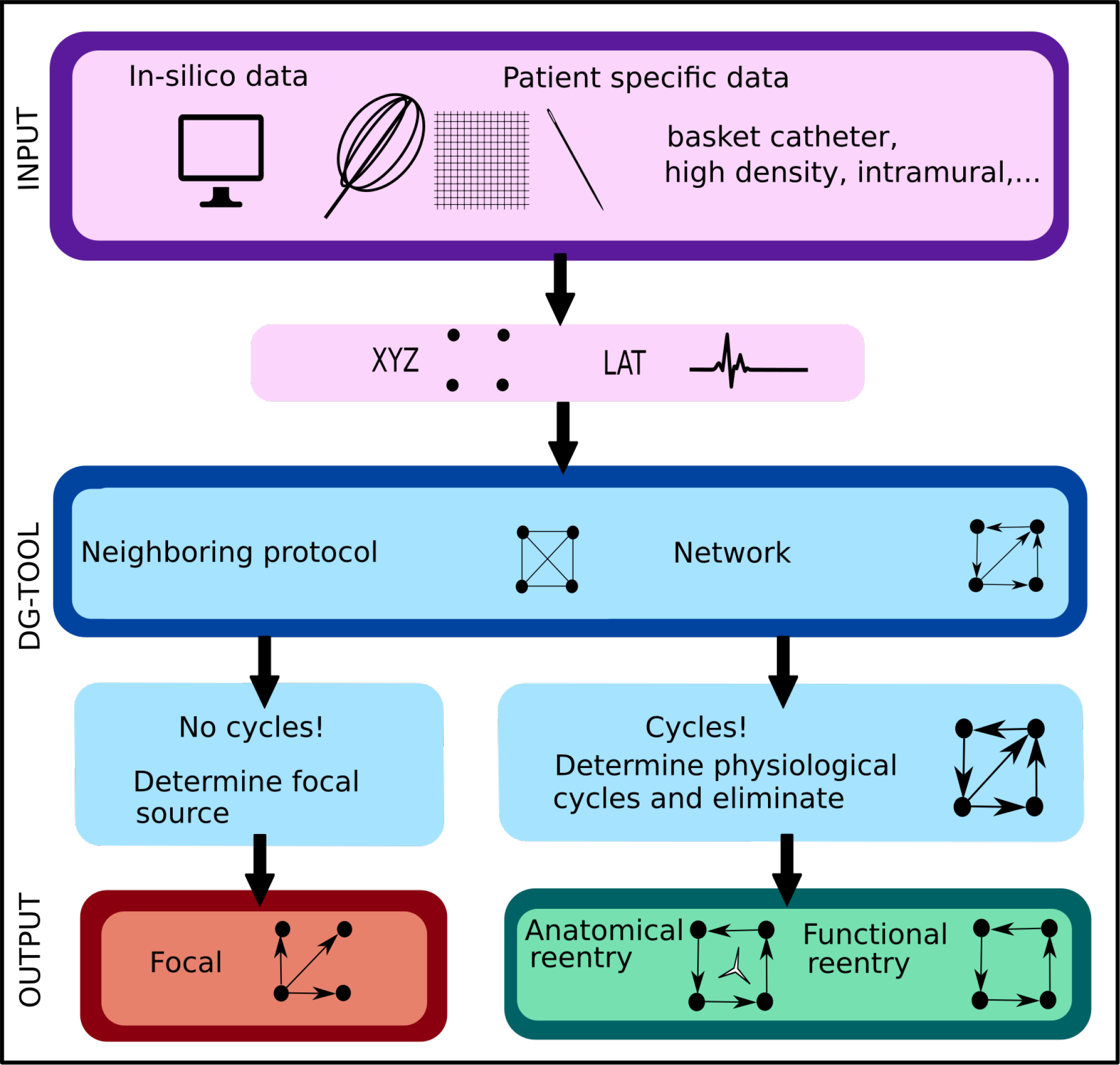
Illustration of the work flow of the DG-mapping tool. As input, the DG-tool requires for a given setup of electrodes the LAT values with the corresponding XYZ coordinates, which can be extracted from either simulation studies or a clinical setup. The input is then processed as follows in the DG-tool. First, for each electrode a neighboring protocol is applied to identify its neighboring electrodes. Second, the network of excitation is generated. Next, we apply a loop-finding algorithm to detect cycles in the network. If cycles are not detected, we locate the source of focal activity. In case cycles were detected, the loops are merged and its center is determined. At the end, the output is visualized. In case of a focal source, arrows pointing away from a (group of) node(s) are shown, while for reentry, arrows will be plotted to visualize the reentry path.

**Fig. 2.**
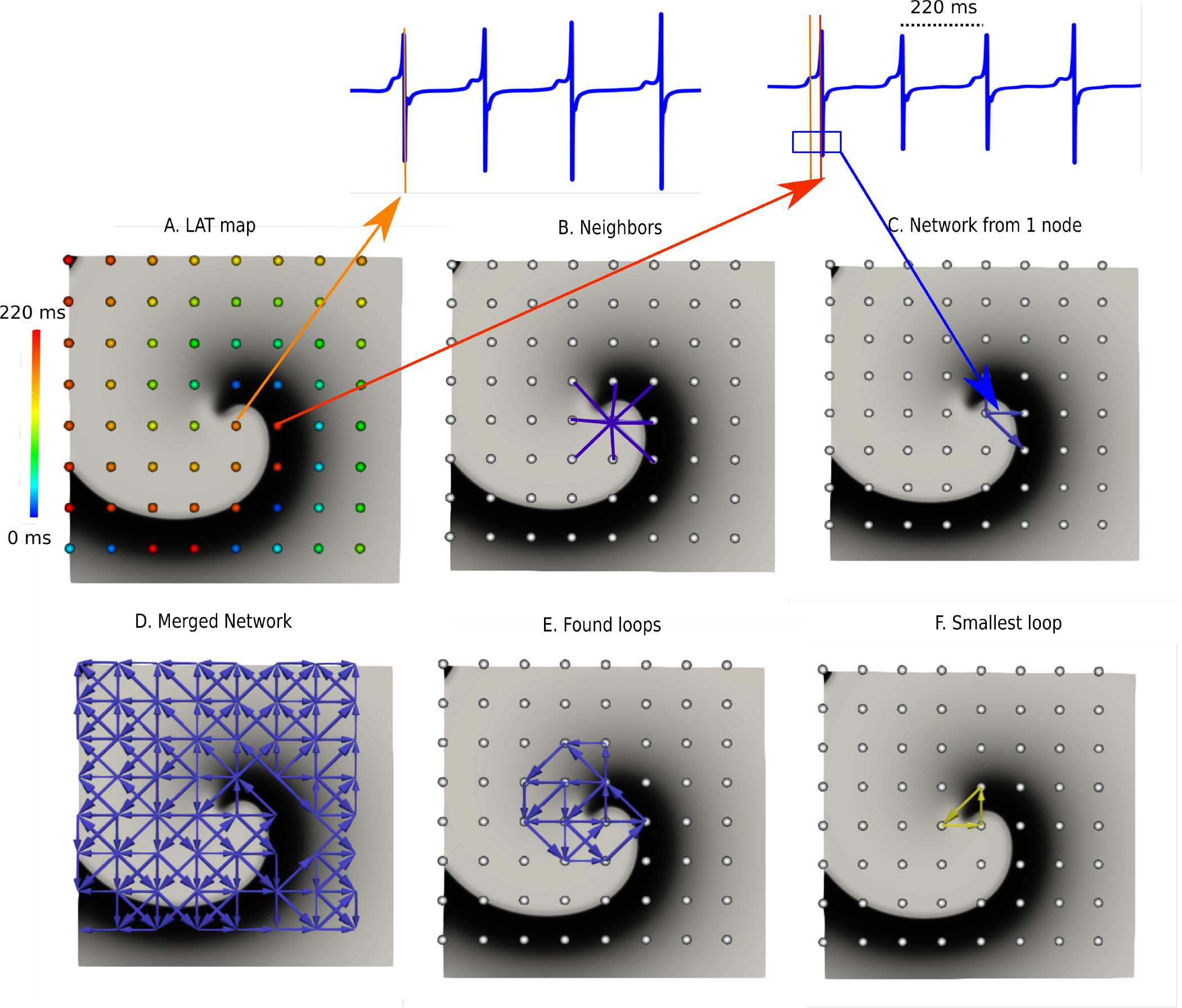
DG-mapping for a simulated rotor with a regular measuring grid. In panel A, the points are colored according to their LAT. Panel B shows an example of the 8 possible neighbors for a single grid point. Panel C represents the corresponding network for the same location. In Panel D the complete network is drawn from which all detected loops in the network are derived, as shown in panel E. The selected (smallest) loop among the bundle is shown in panel F.

**Fig. 3.**
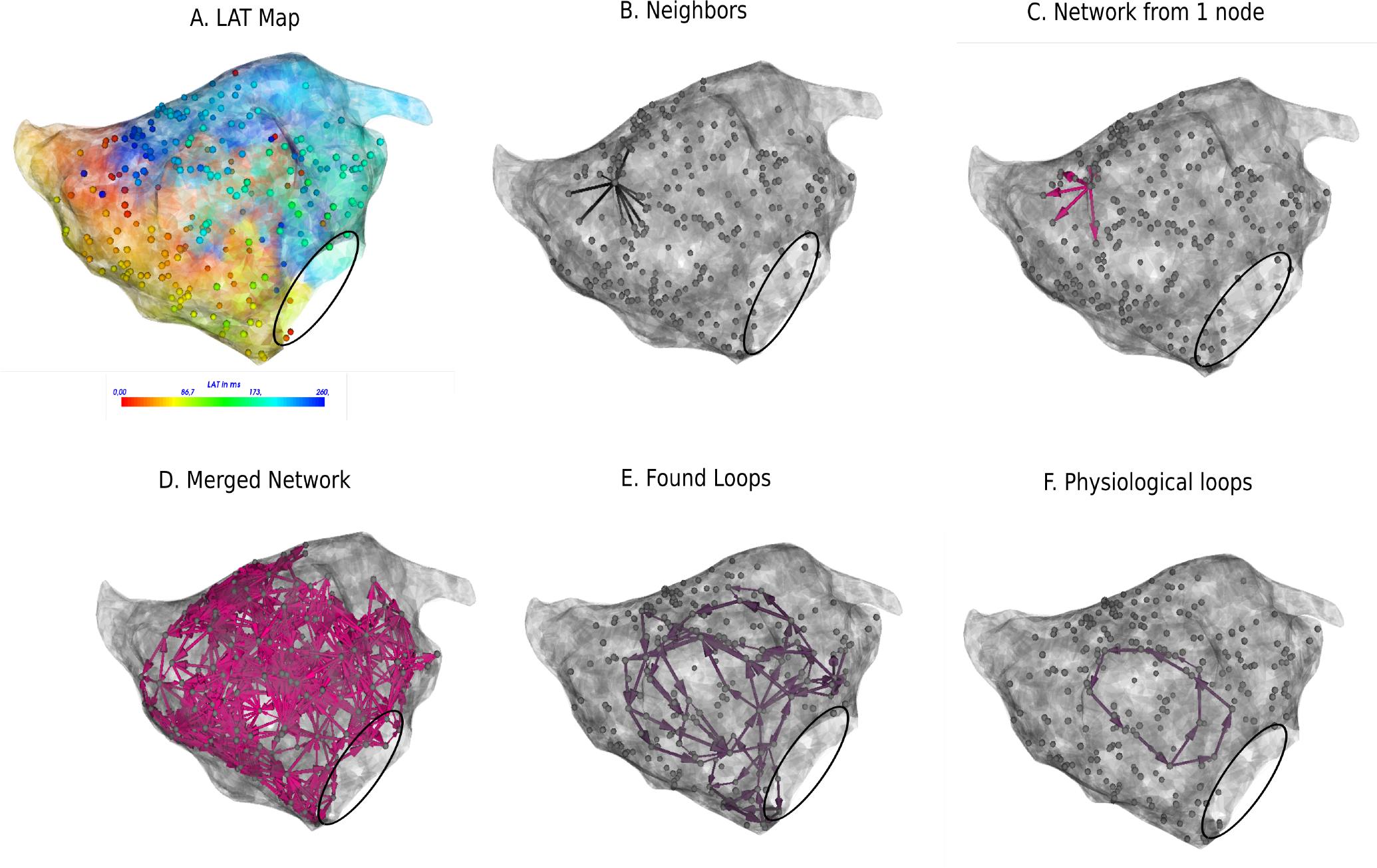
DG-mapping for a regular AT case of localized reentry. The gray dots represent the mapping points, while the black circle denotes the mitral valve. The LAT annotation map in ms is given in panel A. Panel B shows an example of all possible neighbors for a single annotated location. Panel C represents the corresponding network for the same single annotated location. In Panel D the complete network is drawn from which all detected loops in the network are derived, see panel E. The selected loop among the bundle is shown in panel F.

### A Input

The DG-protocol can be applied to a wide range of in-silico, experimental and clinical models of arrhythmia with different types of electrode systems (e.g. basket electrode system, intramural needle electrodes, high density grid data, unstructured electrodes systems…). In the current study, this will be demonstrated with in-silico generated and clinical data.

#### A.1. In-Silico generated datasets

All simulations were performed using the TNNP-computer model for human ventricular cells (15) utilizing the explicit-Euler integration scheme (16) on a GeForce GTX 680 and a GeForce GTX Titan, with single precision. Tthe following different scenarios were simulated: (1) Functional reentry was simulated in 2D (in a domain of 512 by 512 grid points with interspacing of 0.25 mm) and 3D (a simplified model of the human ventricle and an anatomically accurate model of the human ventricle (17)). (2) Anatomical reentry was also simulated in 2D and 3D (anatomically accurate model of the human ventricle). In both scenarios, the S1S2-protocol was applied to obtain rotational activity (18). (3) Focal activity was simulated in 2D and 3D (anatomical model of the human ventricle) by applying 3 stimuli at 3 different locations each 500 ms. All simulations were performed for a duration of 20s.

For each different setup, we implemented either 64 surface electrodes (mimicking 64 electrode-basket catheters (19)), 256 surface electrodes with an interspacing of 0.8 mm (mimicking experimental grid sizes (20)), or 500 intramural electrodes (in the 3D anatomical model) in analogy with the experimental setup by Taccardi et al. (21) In Figure 2, an example of a rotor with 64 electrodes is shown. For these electrodes, we computed local unipolar electrogram (U-EGM) as follows:

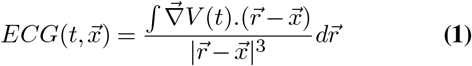

where *t* is time, *x* is the distance to the tissue, *V* is the transmembrane potential, *r* is the integration parameter over the tissue. The XYZ-coordinates of the selected electrodes were also stored for further analysis. The LAT of each electrode was determined by taking the steepest negative slope (−*dV/dt*) of the calculated U-EGM, see also Figure 2A. This coincides with the upstroke of the action potential (i.e., the true moment of activation) (22, 23).

#### A.2. Clinical datasets

Between April and August 2017, 29 patients undergoing ablation of symptomatic ATs at AZ Sint-Jan Bruges Hospital were enrolled in the study. The study was approved by the local ethics committee of AZ Sint-Jan Hospital Bruges. High density (*>* 300 points) endocardial mapping of ATs was performed using a single-electrode mapping and ablation catheter with a distal 3.5 mm tip and three 1 mm ring electrodes (THERMOCOOL SMART-TOUCH Biosense-Webster Inc., Diamond Bar, CA, USA). The bipole of a decapolar coronary sinus (CS) catheter was selected as reference for activation mapping (i.e. peak of CS = 0 ms). The following settings for activation mapping were applied: mapping window set to tachycardia cycle length minus 10 ms and centered at the 0 ms reference, minimum contact force of 4 g, LAT stability of 10 ms, respiratory gating enabled, and color fill calibrated at 5. Bipolar scar threshold was defined at 0.05 mV, and EGMs with bipolar voltages lower than this cutoff were therefore automatically tagged as scar (gray zones) on the activation maps.

Automated and continuous acquisition of points was performed by the CONFIDENSE mapping module (Carto 3 v. 4, Biosense Webster Inc.) using the novel hybrid LAT annotation method (LATHybrid) (24). Each AT case was analyzed offline by DG-mapping after exporting all local activation times (LATs) and the corresponding 3D coordinates. In Figure 3A, an example of the left atrium is shown, with the corresponding LAT map and annotated points.

The tachycardia mechanism was confirmed when ablation resulted in sinus rhythm or in conversion of a second tachycardia.

### B Directed graph mapping protocol

This section explains the DG-mapping algorithms, as shown in the blue panels in Figure 1.

#### B.1. Determine the neighbors in a given system

First, for a given configuration of electrodes, possible neighbors for each electrode are determined. These neighbors cover all possible paths where the wave can travel to, starting from a certain electrode. For regular grids, the neighbors are found by setting a spheric distance around a single point. Hence, a single point incorporates up to 8 neighbors in case of the 2D grid (see Figure 2B), and up to 26 neighbors in case of a regular 3D grid. For an irregular configuration of electrodes, like the clinical AT cases, Delanauy triangulation is applied to determine for each electrode its possible neighbors (see Figure 3B).

#### B.2 Creating network of cardiac excitation

We chose a certain time *t*. Starting from this time, we find *LAT*_1_,.. *LAT*_*n*_ which are the first LAT larger than *t* for each electrode in our system of *n* electrodes. We then draw arrows as follows (25). Suppose electrode 1 and 2 form a pair of neighbors. Assume electrode 1 has *LAT*_1_ and electrode 2 has *LAT*_2_, with *LAT*_2_ *> LAT*_1_, meaning the difference between the two electrodes is *δLAT* = *LAT*_2_ − *LAT*_1_ *>* 0. We allowed a directed link from from electrode 1 to 2 if (25):

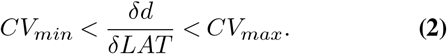

In this equation, *CV*_*min*_, *CV*_*max*_ and *d* represent minimal conduction velocity, maximal conduction velocity and the euclidean distance between the two electrodes respectively. For the simulated examples for ventricular tissue (2D and 3D) we took *CV*_*min*_ = 0.2 mm/ms, and *CV*_*max*_ = 2.00 mm/ms. For the clinical AT cases, *CV*_*min*_ was set at 0.08 mm/ms, according to the lowest physiological conduction velocity in human atria determined by Allessie et al. (26), *CV*_*max*_ was set to maximal 2.0 mm/ms (27). In Figure 2C and Figure 3C, the directed arrows from a single electrode are shown.

Once this first graph was created, a second graph, graph 2, at a time *t* + *δt* was created in exactly the same way as the first graph. We set *δt* = 40 ms. Finally, these two graphs were merged. The resulting graph is the final directed network. For example, in Figure 2D and Figure 3D, the complete network is shown for a simulated case and a regular AT.

#### B.3. Rotational activity

Once the network is created, any type of rotational activity can be found by detecting cycles in the network. A cycle is a closed directed walk without repetition of nodes. In order to find the cycles, a standard breadth-first search algorithm was used. Since the constructed network generally turns out to be rather small and very sparse, this can be done very efficiently. It turns out that detecting all (smallest) cycles through each node can be done almost instantaneously. We ran theoretical simulations on networks with 1 000 000 nodes, and even in these cases all cycles were found in the range of seconds. Clearly, the physical bounds on the number of electrodes that can be placed will be more limiting than the computational work that is needed to process the data. In Figure 2E and Figure 3E the resulting cycles of the network of a simulated rotor and a regular AT case are shown.

In order to find the core of any type of rotational activity, we looked for the smallest cycles in the network, and computed the geometric center. This was performed by grouping all found cycles based on their proximity to the geometric center. If the centers lie closer to each other than a specified threshold, the cycles were considered to belong to the same core. Afterwards there was an optional pass which merges bundles of cycles if they shared nodes. Finally, the centers of each bundle were defined as the core of rotational activity. In Figure 2F, and Figure 3F, the selected cycles are shown.

#### B.4. Focal activity

Focal activity was detected as a source node, i.e., a node which has a non-zero out-degree, and an in-degree equal to 0. These can be found immediately by doing a single pass over all nodes. Then, the LATs were bundled in certain intervals, to reduce the inter-variability in the LAT values. Afterwards, we reconstructed the network with these bundled values. We then checked if regions with only outgoing arrows were present. The middle of these regions corresponds to the source of the focal activity.

### C Additional features of network theory

In addition to finding rotational and focal activity, we derived additional properties of the network.

#### C.1. Region of influence

For each network containing rotors, we can determine a ‘region of cycles’ and a ‘region of influence’. The region of cycles contains all nodes (electrodes) which are part of cycles for a particular rotor. Second, for each non-marked point we can determine the closest ‘region of cycles’ in terms of network arrival time distance and relate it to that region. As a result, for each point we can determine which source excited it. This is called the ‘region of influence’.

In order to construct the region of influence the following algorithm was implemented. For a given network, all *n* cores were determined, *c*_1_,…,*c*_*n*_. For each core, we first determined all nodes which are part of cycles of the network (*C*_1_,…,*C*_*N*_), i.e. the regions of cycles. Then, each node was added to the core *c*_*i*_ to which it had the shortest path to one of its nodes in *C*_*i*_. In this way, each core is assigned a region of influence.

#### C.2. Wave averaging

Another application of the constructed network is wave averaging to interpret the cardiac excitation pattern. In general, wave averaging consists of decomposing any directed link into smaller arrows which are each projected on a given 3D geometry, followed by averaging all arrows within 1 cm of a node. In more detail, the following steps are taken in the wave averaging algorithm. First, each LAT-node was projected on the mesh. Second, each arrow of the directed network was also projected on the mesh in the following way: (A) Each arrow 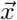 between two LAT-nodes on the mesh was divided in two arrows, both projected on the mesh, 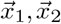. (B). Each of these arrows 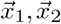 is then again divided into two arrows and projected 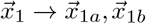 and 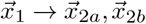. Therefore, one arrow 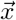, was divided in 4 projected arrows on the mesh 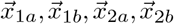. Third, for each node *n* on the mesh, each directed arrow starting from this node as well as each connection of each node within 1 cm from the original node *n* was averaged. The collection of these averages was then plotted on the mesh.

### D Phase mapping protocol

LAT values were used to construct the excitation patterns in phase-space. First, a sawtooth wave, with amplitude ranging from −*π* to *π* is constructed based on these LAT values. Afterwards, values are adapted with their 2*π* equivalent within the range of −*π* to *π* in phase-space. Next, in both x and y direction, the phases were derived and a linear combination with the Sobel (we also tested the Nabla) kernels to detect the singularities was applied. This protocol was previously presented (28–32), but instead of the Hilbert transform we made use of the sawtooth wave for the phases. In 3D, the heart was sliced in 3 orthogonal directions and the protocol was applied on each slice. However, as the shape of the ventricle model is complex, only grid points with complete circumference in the heart were taken into account, so convolution did not result in false positives on the edges (33). However, this did not result in detection of the filament of the rotor as the density (500 intramural points) was too sparse. We therefore calculated the phase singularities on the surface of the tissue and detected eventually the phase singularities of the spiral in 3D. A binary detection threshold was applied to the convolution (32), set to 95% of the maximal detected value in phase-space.

### E Introducing LAT variation

In the clinical setup, identification of LAT either by automated algorithms or manual annotation by operators can vary due to several factors such as accuracy of the detection algorithms, operator experience, signal quality and noise (34). Therefore, we included LAT variation in our analysis, and compared the accuracy of DG-mapping with phase mapping. In order to obtain LAT variation in the simulations of functional reentry, random Gaussian noise was added with standard deviations *σ* = 5, 10, 15, 20, 25, 30 ms in the simulation of functional reentry with a configuration of 64 and 256 electrodes. We divided the activation times obtained during a simulation in 25 different frames with 520 ms separation to exclude any overlap in activation times. For each frame, we randomly added Gaussian noise 1000 times, so in total, we compared 25000 different frames per LAT variation *σ*. The center of the rotor was detected through DG-mapping and phase mapping. For DG-mapping, the geometric center of all cycles belonging to the same core was computed. Afterwards, the median value was taken as the true center of the rotor. In addition, only the center with the highest number of cycles was taken into account. We classified the outcome as correct if only one single core was found within 1 cm of the true core. The incorrect diagnosis was classified in 3 different types: incorrect cores (i.e. cores outside a 1 cm radius of the true core) in combination with the correct core (error type 1), only incorrect cores (error type 2), or no cores (error type 3). For the percentage correct diagnosis *p*, we computed a 95% confidence interval via *p* ± 1.96*SE* where *SE* is obtained from a robust sandwich estimator (35, 36) that accounts for the correlation structure (i.e. the 1000 replicates within one time frame are expected to be correlated). In the supplementary material, we also simulated noise from the skewed log-normal distribution to study the robustness of the methods for different types of noise distributions. In addition, we also presented the outcome as a function of the distance from the true core (instead of taking 1.0 cm).

## 3. Results

### A In-silico models of functional and anatomical reentry and focal activity

The accuracy of DG-mapping was tested in different in-silico models as described in the methods section. First, for functional reentry (see Figure 4A), we simulated a 2D rotor with a configuration of 64 electrodes (A1) and 256 electrodes (A2). In 3D, functional reentry was induced in a simplified model of the ventricles with 64 surface electrodes (A3), and in an anatomical model of the ventricles with 500 intramural electrodes (A4). In all four setups, DG-mapping was able to accurately detect functional reentry and correctly determine the location of the core of the rotor for the entire length of the simulation (20s duration). The smallest cycle and corresponding core are shown in yellow for each setup. Second, DG-mapping was validated in two models of anatomical reentry (Figure 4B): a 2D anatomical circuit with 64 electrodes (B1) and a 3D anatomical reentry with 500 intramural points in the model of the ventricles (B2). In both models, DG-mapping correctly identified the reentrant path around the obstacles for the entire length of the simulation (20s). The shortest reentry loops are again depicted in yellow. Third, focal activity was simulated in 2D (64 electrodes) and 3D (500 intramural electrodes) (Figure 4C), by repetitively stimulating 3 different locations. Again, DG-mapping identified the electrodes most closely to the site of stimulation, see C1 (yellow arrows) and C2 (black circles).

**Fig. 4.**
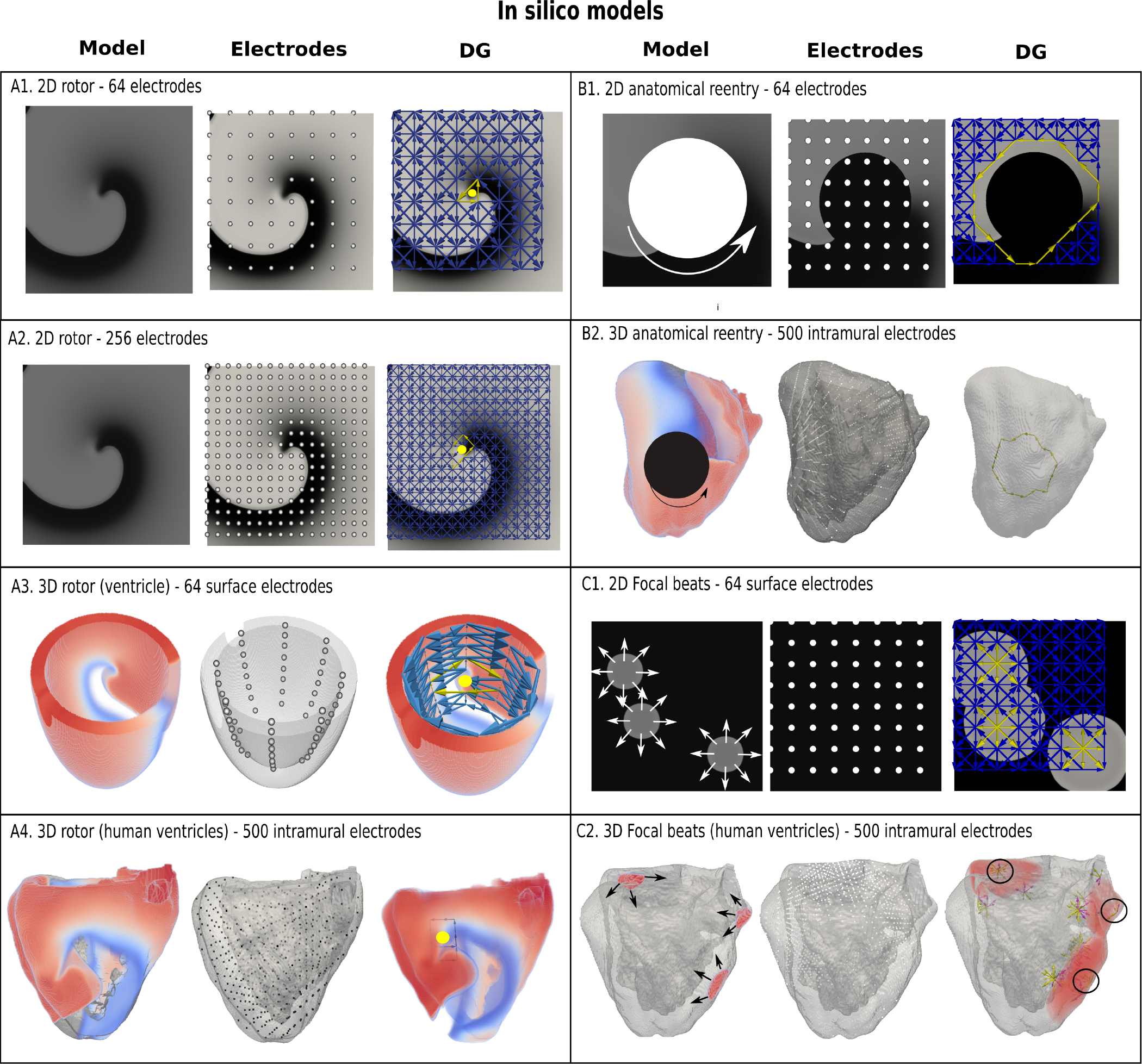
Examples of DG-mapping for different in-silico models. Model A1 and A2 show a rotor in 2D with 64 and 256 electrodes. Model A3 shows a rotor in a simplified model of the ventricle with 64 surface electrodes. Model A4 depicts a rotor in an anatomical model of the ventricles with 500 intramural electrodes. The smallest cycles and their geometrical centre are indicated in yellow. Model B1 depicts anatomical reentry in 2D with 64 electrodes and Model B2 in a model of the human ventricles with 500 intramural electrodes. The smallest reentry loop around the obstacle is again depicted in yellow. Model C1 represents 3 focal sources in 2D with 64 electrodes, whereby DG mapping shows the sources in yellow. Similar results with 3 focal beats in the ventricles are shown in model C2, indicated with 3 black circles.

### B Clinical dataset

To establish proof of concept in the clinical setting, we retrospectively and blindly analyzed 31 cases of regular atrial tachycardia (AT). For clarity, in Figure 3, all the steps of the DG-mapping protocol were demonstrated on an AT case of a localized reentry. In general, the atria have a complex structure. In case of reentry during AT, the electrical waves circle around obstacles such as the valves, the veins or scar tissue, creating a (sustained) reentry loop. Ablation aims to terminate the reentry loop so that the circular electrical conduction can no longer be sustained.Therefore, it is important to precisely determine the location of the activation pathway. The accuracy of DG-mapping was compared to the standard diagnosis i.e. type of arrhythmia and location of the circuit/focal activity as determined by the electrophysiologist (EP) based on the activation map and the ablation result. The overall results are summarized in Figure 5. Out of 31 cases, 20 were due to macro-reentry, 6 due to localized reentry and 5 due to focal activity, see also the supplementary Table S1.

**Fig. 5.**
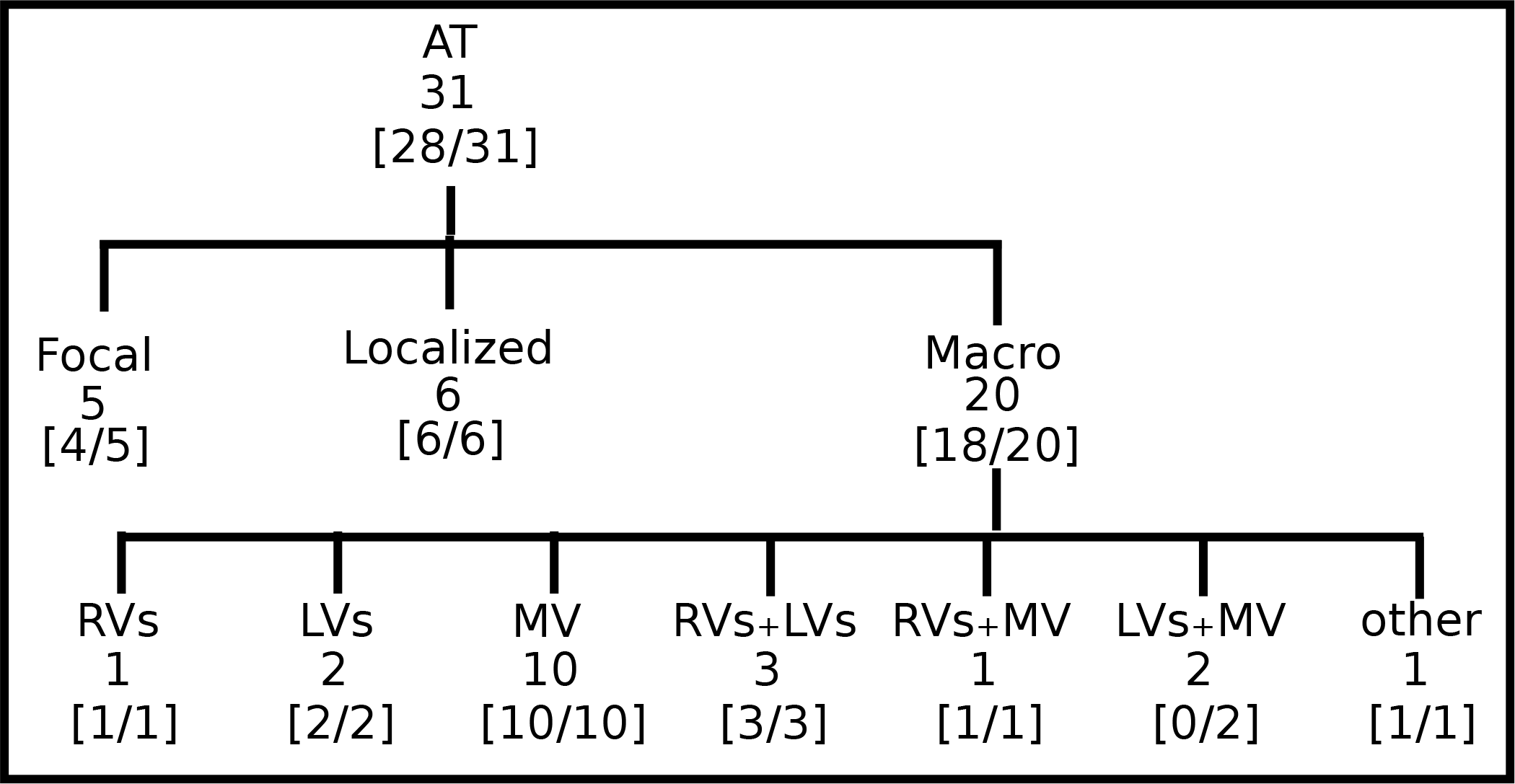
Accuracy of DG-mapping in clinical AT. The gold standard was compatible with focal source (5), localized reentry (6) and macro-reentry (20). Macro reentry was categorized in reentry around the right veins (RV), the left veins (LV), mitral valve (MV), around RV + LV, around RV + MV, around LV + MV, and other types of reentry (e.g. in the right atrium). In brackets the accuracy of DG-mapping is given.

In 9 cases with reentry, the operator was not sure about the reentry mechanism purely based on the LAT activation map, formulating several hypotheses. The gold standard was taken as the diagnosis matching the ablation endpoint. Compared to this gold standard diagnosis, DG-mapping identified the exact same mechanism and location in 28 out of 31 cases (90.3%, 95% exact binomial confidence interval 74.2% − 98%). In 3 out of 31 cases, the diagnosis of DG-mapping did not fully match with the gold standard. In 2 cases of double loop reentry, (case 6 and 14), DG-mapping identified only one single loop. In the other case (case 22), the mapping data indicated focal tachycardia, whereas DG-mapping identified localized reentry at the same location. However, in all 3 cases, DG-mapping would have pointed to the correct ablation target, meaning that DG mapping correctly identified the ablation target in 31/31 cases.

Representative cases are shown in Figure 6. Panel A depicts a macro-reentrant AT around the right veins conducting over the roof. Ablation of the roof resulted in prompt termination of the AT. Blinded analysis by DG-mapping revealed a selected loop at the same location (middle panel). Panel B shows a localized reentry at the anterior wall, rotating around local scar tissue. Ablation from the scar to the mitral valve terminated AT. DG-mapping (middle) as well as wave averaging (bottom) identified the same location of the localized reentry. In panel C, activation mapping and ablation was conform with focal tachycardia at the septum. DG-mapping (in the absence of loops) pointed to focal activity as well (middle panel). We also tested the wave algorithm for each case, as shown in the bottom panels of Figure 6. Wave averaging was compatible with macro reentry (A), localized reentry (B) and focal activation (C).

**Fig. 6.**
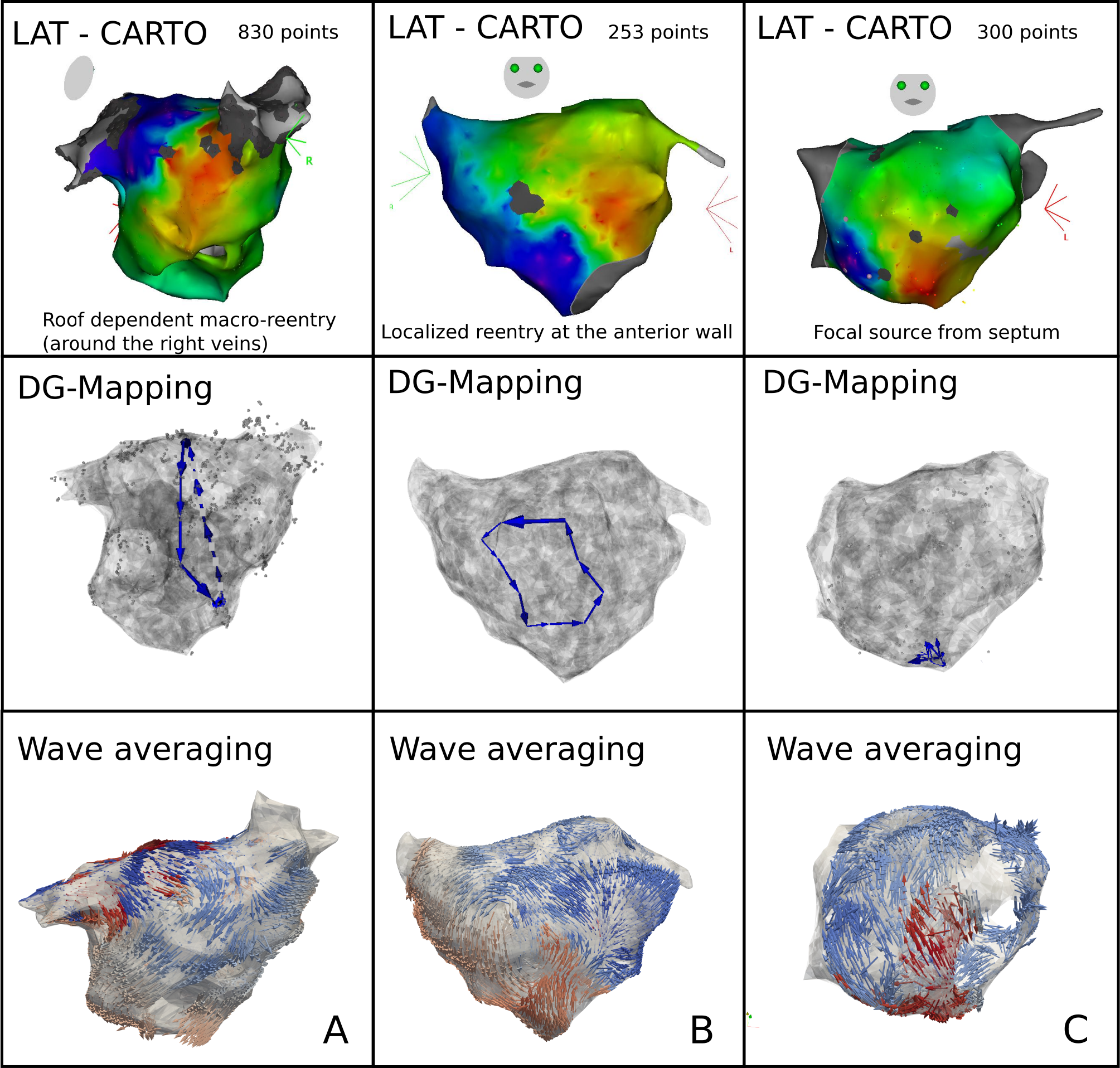
Representative examples of DG-mapping in 3 cases of clinical AT: macro-reentry (A), localized reentry (B) and a focal source (C). For each case, the activation map and the corresponding DG-map and network based wave averaging is given.

### C Comparison with phase mapping under LAT-variation

In the clinical setup, LAT’s can vary due to several factors such as accuracy of the detection algorithms, operator experience, signal quality and noise (34). Therefore, the performance of DG-mapping was compared to phase mapping in the model of a single rotor with 256 electrodes, now by adding Gaussian white noise to the LAT’s (Figure 7, upper panels). Overall, we observed that DG-mapping retained its accuracy to detect rotors at increasing noise levels, whereas phase mapping becomes less precise (middle panel of Figure 7): for small variation levels (5 ms), DG-mapping is 100% accurate, while the accuracy of phase mapping decreased to 74.17%. For 15 ms, phase mapping became highly unreliable (accuracy of 30.22%), while DG-mapping had an accuracy of 95.49%. For 20 ms, this difference was even more pronounced: DG-mapping maintained an accuracy of 81.19% while the accuracy of phase mapping dropped to 1.08% (p-value *<* 0.0001). Moreover, in case of incorrect diagnosis (lower panels), phase mapping detected extra false cores (type 1 error) whereas in the DG method incorrect diagnosis was due to no detection of the the core (type 3 error). Noise analysis was repeated for the 2D model with 64 electrodes. A similar trend was found with DG-mapping being more accurate (91%, 83%, 68%, for a noise level of 10, 15 and 20 ms) versus phase mapping (75%, 51%, 21% respectively). All of these differences were highly significant (p-value *<* 0.0001), see also Table S2 in the supplementary material. We also varied the distance to the true core for which we retained a diagnosis as correct, as shown in Figure S2. As explained in the supplementary material, due the discrete nature of phase mapping, we can only compare phase mapping and DG-mapping above a certain threshold. In these cases, DG-mapping always exceeded phase mapping. We also tested the effect of the underlying distribution of the LAT values. Modeling LAT variation with the skewed lognormal distribution did not alter the conclusions of the results, see also Figure S1. Finally, to test the specificity of DG-mapping, we applied DG-mapping in a point simulation model without functional reentry (256 electrodes, LAT variation ranging from 0 ms to 30 ms). In these cases, DG-mapping never identified any rotors, resulting in a specificity of 100

**Fig. 7.**
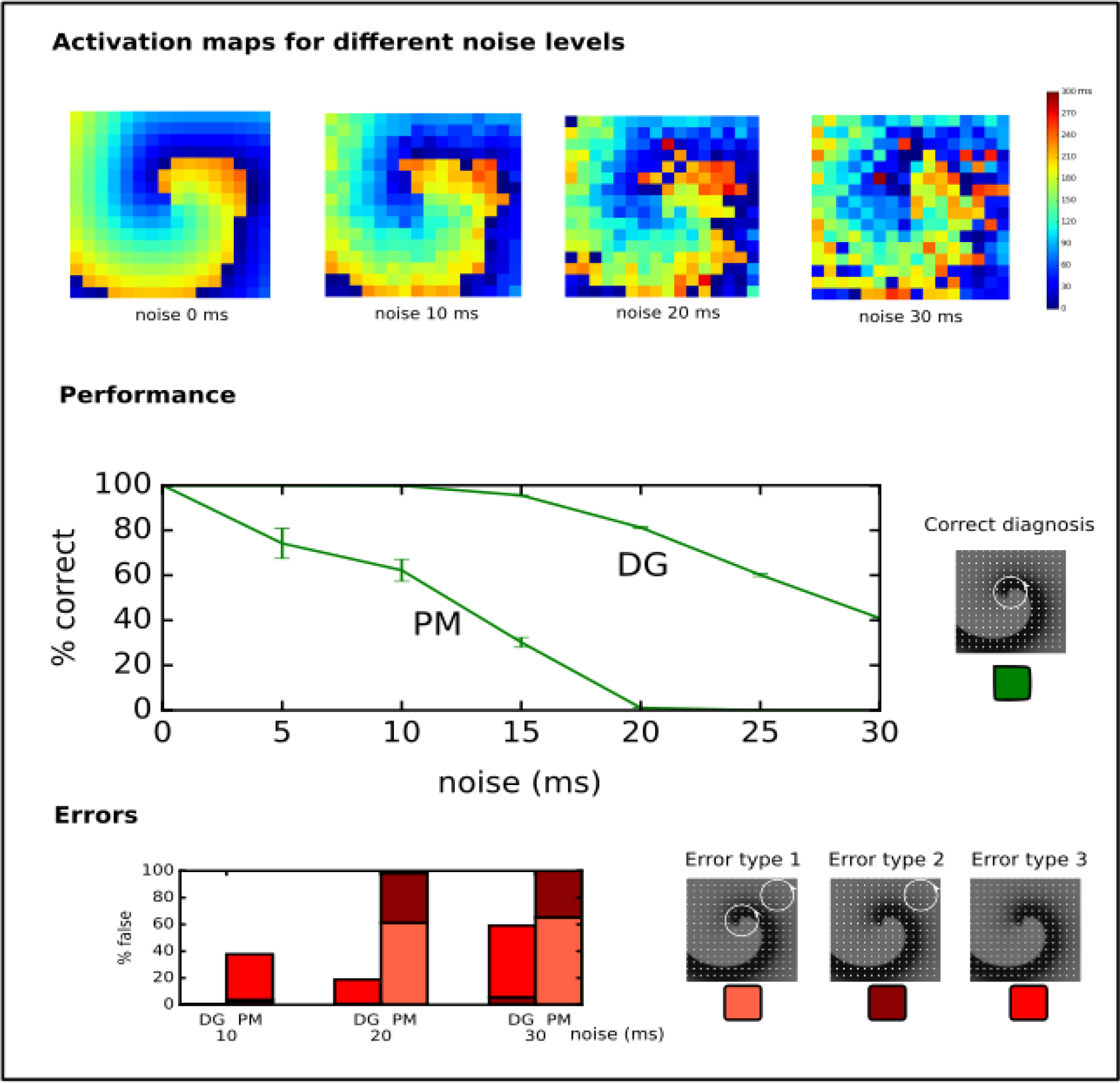
Performance of DG-mapping and phase mapping (PM) while adding Gaussian noise with an increasing standard deviation (ranging from 0 to 30 ms) in a single 2D rotor model with 256 electrodes. The upper panels show representative activation maps for different levels of noise. The middle panel shows the performance of DG-mapping versus phase mapping for these different noise level. The bottom panel shows the type of errors in case of failure for both methods. Error type 1 is a detection of a false core in addition to the correct core, error type 2 is only a detection of a false core and error type 3 is no detection of cores at all.

### D Region of influence

Describing cardiac excitation as a network, allows to extract additional information. Besides wave averaging, DG-mapping allows to identify the spatial region which is excited by a certain source. In case of normal excitation, a single source (sinus node) excites the whole medium. However, in case of an arrhythmia with multiple sources, each source excites a given region, which we call the region of influence. We hypothesized that DG-mapping, by containing complete spatio-temporal information, can determine this area of influence. This concept was evaluated in 2 different setups (see Figure 8A). We determined the region which contained the electrodes belonging to cycles (region of cycles), as well as the region of influence. Obviously, for a single rotor, the region of influence spans the entire set of electrodes. For 4 different rotors, one can observe an area of influence for each given source.

**Fig. 8.**
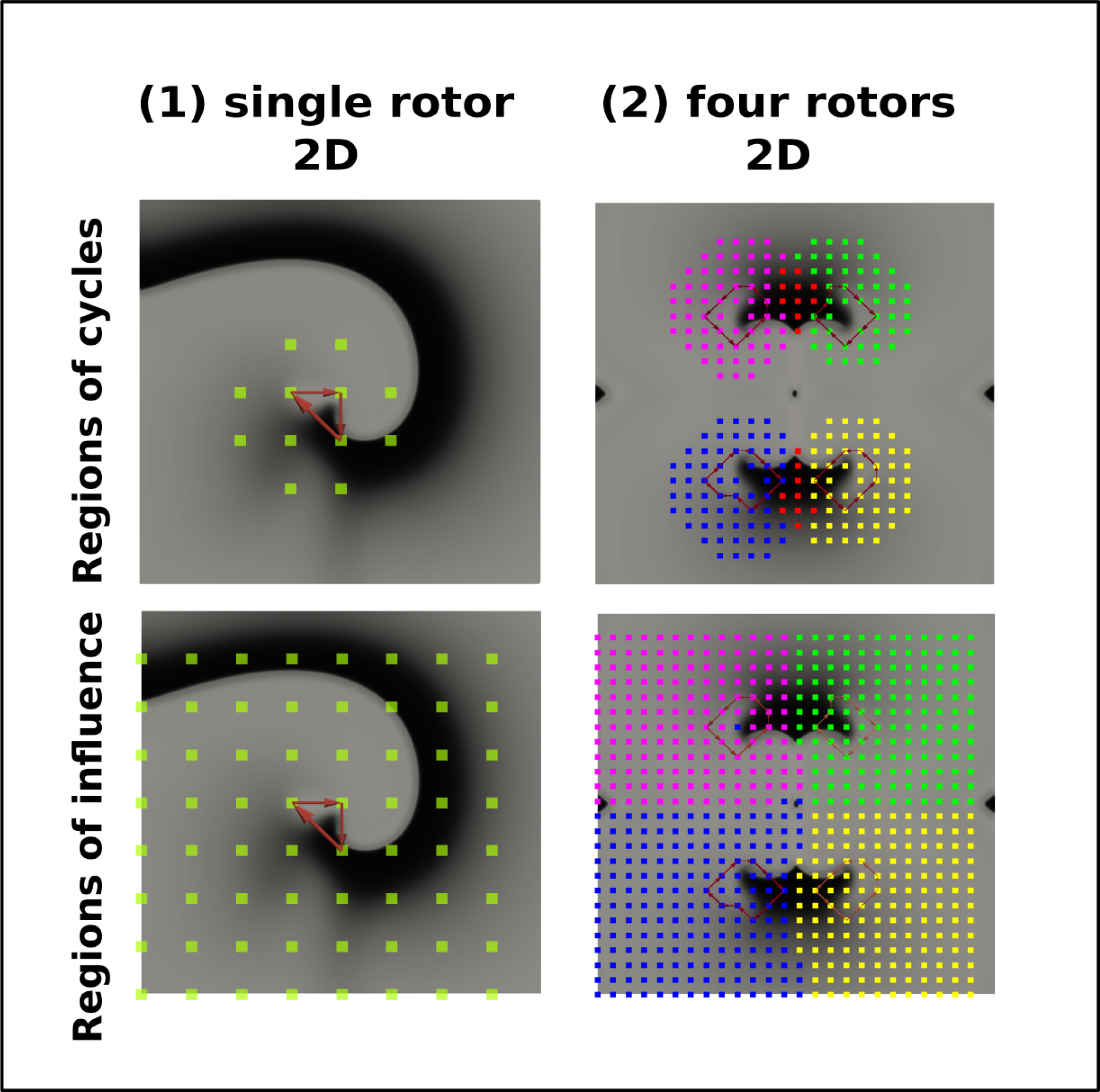
The regions of cycles and the regions of influences for model A (single rotor) and model B (four rotors). Model A and B were simulated with 64 resp. 625 electrodes. The smallest cycles are depicted in red. Different regions of cycles were depicted in different colors, while the overlap is shown in red. Similarly, different regions of influence are shown in different colors, dividing model B into four different regions.

## 4. Discussion

### A Main findings

In this paper, we demonstrated that a directed network can be used to describe electrical activity in the heart in order to find the mechanism of the arrhythmia. This novel method is robust, fast, general, accurate and can be applied to a wide range of in-silico, experimental and clinical models of arrhythmias. First, we showed that DG-mapping can find functional reentry, anatomical reentry and focal activity in in-silico ventricular models of the heart (see Figure 4). We tested using intramural electrodes, 64-basket electrodes and regular grids with different numbers of electrodes (64-256). Second, we tested DG-mapping om 31 clinical cases of regular AT, see Figure 6. Compared to the gold standard, DG-mapping identified the exact same mechanism and location in 28 out of 31 cases, whereas it identified the correct ablation target in all 31 cases. These results suggest that DG-mapping could potentially lead to an improved treatment of cardiac arrhythmia.

### B Network theory

To our knowledge, so far only limited research has focused on network theory to understand cardiac arrhythmias. In the study by Zahid et al. (37), undirected networks were used to find the minimal number of nodes which need to be ablated to separate two regions in the heart. This region was then proposed as the optimal ablation site. In the study by Tao et al. (38), the authors showed that ablation of persistent AF is associated with improvement in both local and global connectivity within the communication networks. However, in neither of the above studies, excitation was interpreted as a *directed* network. Zeemering et al. (39), applied a directed network to describe AF, by accumulating multiple time frames. However, in contrast with DG-mapping, this methodology precluded the possibility of detecting rotational activity, and does not represent the actual wave excitation. In a model of chronic atrial ventricular block, we used directed networks for the first time to determine the mechanism underlying Torsade de Pointes (25). However, again, we did not use it to fully describe the electrical excitation as in this work. Therefore, to our knowledge, this is the first study where directed networks were used to describe electrical excitation, building on our previous work in the CAVB dog (25).

### C Advantages of DG-mapping

First, we showed that DG-mapping could be used to reliably detect rotational activity even after adding LAT variation. For instance, in the model presented in Figure 7, we found that in case of 15 ms standard deviation of noise, phase mapping was only 30% accurate, while DG-mapping was still 96% accurate. This difference in accuracy can be explained by the holistic nature of DG-mapping. Even in the presence of a number of electrodes with a wrong LAT annotation, DG-mapping can still identify the correct location of the rotor based upon the other electrodes. In addition, DG-mapping also takes into account the number of cycles which are found for each rotor (with a higher number of loops indicating a higher likelihood of an actual rotation). In contrast, phase mapping finds phase singularities locally. Therefore, a misplaced LAT can easily give rise to false positives/negatives, potentially resulting in incorrect clinical decisions.

Second, DG-mapping automatically detects anatomical reentry and focal activity, a feature which phase mapping lacks. Very recently, another methodology was described by Oesterlein et al. which can also automatically detect anatomical reentry (40). The approach uses a different methodology based on integral measures: determination of activated area and its relation to the cycle length of the arrhythmia, while our DG-mapping directly analyses the local propagation of the excitation wave. It would be interesting to compare this method with DG-mapping especially in clinical settings.

Third, DG-mapping offers additional features which can be derived from the directed network. DG-mapping can determine all electrodes belonging to any cycle which are part of the same rotational activity (see Figure 8A), and detect for each rotational core its region of influence (Figure 8B). This offers the possibility to detect all electrodes activated by a specific rotational activity, and could detect the dominant driving source of the arrhythmia as a primary target of ablation. Also, the wave averaging technique allows to create maps of the wave propagation, which can provide additional guidance during catheter ablation.

Finally, another advantage is that DG-mapping is universal. It can be applied to any type of recording system, with varying number of electrodes, the inter-electrode distance or site of recording, as shown in Figure 4.

### D Clinical implications

As shown in this paper, DG-mapping can be of added value in the ablation of regular AT. Despite improvements in activation mapping (RHYTHMIA by Boston Scientific, Coherent mapping system by Biosense Webster), interpretation of activation maps remains challenging and operator dependent (12, 41). We demonstrated that DG-mapping automatically identified the same mechanism as the electrophysiologist (EP) in 28/31 cases of regular AT, but did find the correct ablation target in 31/31 cases. Moreover in 9 cases with reentry, the operator was not sure about the mechanism based on the LAT activation map and formulated several hypotheses (see supplementary Table S1). Therefore, DG-mapping could aid physicians in finding the correct diagnosis according to the ablation target. Currently, in case of doubt, the operator can perform entrainment mapping whereby the post pacing interval (PPI) is compared to the tachycardia cycle length (TCL) to localize or confirm the correct reentry circuit (42). Furthermore, DG-mapping also automatically detects focal activity and its location, making it a complete diagnostic tool for AT. The wave averaging technique allows to create maps of the wave propagation, which can provide additional guidance during catheter ablation. Another advantage is that DG-mapping is instantaneous, and could therefore fasten the ablation procedure.

Another important potential application could be AF. AF is often referred to as the most common arrhythmia in clinical practice with an estimated prevalence of 2% and is associated with a five-fold and two-fold higher risk of stroke and death, respectively (43). Catheter ablation of AF yields moderate success rates (44–46), which is related to the lack of understanding of AF mechanisms. Different mechanisms for AF have been described such as focal activation, dissociated activity or stable rotors (47–52). Currently, both researchers and electrophysiologists rely on activation mapping or phase mapping for the analysis of AF. Recently, initial studies suggested good outcomes after ablation of rotors guided by phase mapping (19, 53). However, new studies emerged contradicting this study (54–56). It was shown that phase mapping easily generates false positives (57, 58), especially due to LAT variations on the signals and large inter-electrode distances. In our study, phase mapping showed similar results when adding LAT variations, whereas DG-mapping maintained a high accuracy. It remains to be seen whether DG-mapping will offer new insight in AF mechanisms, however, the holistic nature of the method (as explained in the advantages of DG-mapping) might overcome the problems with phase mapping, as currently used in the clinic.

In conclusion, translating cardiac arrhythmias into directed networks as described in the current work offers the beginning of a new area of research. There exists a whole range of different algorithms in network theory (e.g. edge density, centrality measures, etc. (59–62)), which can possibly be applied to the constructed networks to increase our understanding of cardiac arrhythmias.

### E Limitations

As this paper was a proof-of-concept, many different settings are not yet tested. For example, it remains to be tested how DG-mapping will characterize cardiac excitation in more complicated settings with multiple meandering rotors, including wavebreaks, or in complex fibrotic tissue. A limitation of DG-mapping is that it requires at least one full cycle (as DG-mapping can only find closed loops) of a circular rotation, while phase mapping finds phase singularities instantaneously. In the clinical setting, it remains to be seen if DG-mapping can advance the understanding of AF, VT and VF. For these cases, DG-mapping requires the arrhythmia to be mapped first, which is not always possible (e.g. due to hemodynamic instability in VT/VF etc.). Future studies are needed to further evaluate DG-mapping in different types of arrhythmias.

## 6. Supplementary material

**Table S1.** Summary of the 31 clinical cases of regular AT. For cases 1-16, we used non-interpolated data given by the CARTO system. For cases 17-32, the interpolated data from CARTO was used. The columns represent the case number (#), the database code, the number of measured points (# el), the cycle length (CL), the mechanism found by the EP and the mechanism found by DG-mapping. We put a ✓if the mechanism fully agrees, and a ☑ if the ablation point agrees, but a different mechanism is seen. SCC = slow conductive channel, PV = pulmonary vain, RSPV = right superior pulmonary vain, RV = right vain, LAA = Left Anterior Appendage.

| # | code | # el | CL | Mechanism LAT map CARTO (EP) | Mechanism DG |
| --- | --- | --- | --- | --- | --- |
| 1 | ATM1 | 865 | 250 | **Macro reentry:** perimitral circuit clockwise | ✓ |
| 2 | ATM2 | 784 | 300 | Hypothesis 1: Large macro reentrant circuit sticking close to the right veins, traveling up on the anterior wall, over the roof and down on the posterior wall. There is a SCC on the anterior RSPV side.<br>Hypothesis 2: **localized reentry** circuit at the anterior wall next to the RSPV through the SCC ablation was done on the SCC, which agrees with both hypothesis. | ✓ (hypothesis 2) |
| 3 | ATM3 | 219 | 450 | Hypothesis 1: Large **macro reentry:** roof traveling up on the anterior wall, over the roof and down on the posterior wall.<br>Hypothesis 2: localized reentry on the anterior wall. | ✓ (hypothesis 1) |
| 4 | ATM4 | 1722 | 225 | Double loop: **macro reentry:** perimitral reentry counterclockwise and **macro reentry** around the LV over the roof. | ✓ |
| 5 | ATM5 | 1567 | 200 | **Macro reentry:** perimitral reentry clockwise | ✓ |
| 6 | ATM6 | 1752 | 214 | Hypothesis 1: double loop: **macro reentry:** perimitral counterclockwise and **macro reentry** around the LV over the roof.<br>Hypothesis 2: roof dependent macro reentry, with a pseudo (slower) reentry around the mitral valve. | Only perimitral counterclockwise reentry found but the ablation target would be at the same location ☑ |
| 7 | PC1 | 1753 | 245 | **Focal** | ✓ (no loops detected) |
| 8 | PC2 | 889 | 290 | **Macro reentry:** roof | ✓ |
| 9 | PC3 | 896 | 302 | **Macro reentry:** perimitral reentry clockwise | ✓ |
| 10 | PC4 | 431 | 240 | Perimitral circuit counterclockwise. This was ablated and the AT changed and switched through the roof with a CL of 228 ms. However, retrospective offline analysis revealed a localized reentry below the left atrial appendage (at the base of the LAA). Original ablation points agree with localized reentry. | ✓ (localized-reentry) |
| 11 | PC5 | 847 | 250 | **Macro reentry:** peritricuspid reentry in the right atrium | ✓ |
| 12 | PC6 | 444 | 278 | **Macro reentry:** mitral reentry counterclockwise | ✓ |
| 13 | PC7 | 354 | 340 | **Macro reentry:** perimitral circuit | ✓ |
| 14 | PC8 | 938 | 260 | **Macro reentry:** perimitral circuit counterclockwise and a second loop going through the roof. The two loops are merging at the left veins. | Only perimitral reentry found but the ablation target would be at the same location ☑ |
| 15 | PC9 | 283 | 304 | **Focal** | ✓ (no loops detected) |
| 16 | PC10 | 3524 | 240 | The online LAT map was uninterpretable, however roof line ablation was performed. Offline retrospective re-evaluation showed a possible atypical focal activity from the scar on the roof. The ablation roof line went through that scar and that possible terminated the AT. | ✓ (no loops detected) |
| 17 | CD201 | 475 | 260 | Localized **localized reentry** at the crosspoint at the appendage and the roof, anterior superior appendage from the left atrium. | ✓ |
| 18 | CD203 | 1092 | 240 | Localized **localized reentry** at the anterior wall near the mitral annulus at 11 o' clock. | ✓ |
| 19 | CD204 | 1091 | 240 | Hypothesis 1: **macro reentry**: perimitral circuit counterclockwise using a SCC in between the mitral annulus at 10 o'clock and a scar at the anterior wall.<br>Hypothesis 2: the same loop as in hypothesis 1, but a second loop is found through the same SCC turning around the scar at the anterior wall.<br>Ablation at the SCC terminated the AT (according to hp1). | ✓ (hypothesis 1) |
| 20 | CD205 | 463 | 290 | Hypothesis 1: **macro reentry**: perimitral circuit clockwise using a SCC in between the mitral annulus at 10 o'clock and a scar at the anterior wall.<br>Hypothesis 2: the same loop as in hypothesis 1, but a second loop through the same SCC turning around the scar at the anterior wall counterclockwise. (the size of this second loop cannot be determined).<br>Ablation at the SCC terminated the AT (according to hypothesis 1) | ✓ (hypothesis 1). |
| 21 | CD206a | 612 | 250 | **Focal** breakthrough at the septum inferior and right inferior PV. | ✓ (no loops detected) |
| 22 | CD206b | 360 | 250 | **Focal** AT at posterior septum superior, but 2 signals were not able to be explained by the focal activity.<br>After diagnosis from DG-mapping, the map was reconsidered and localized reentry agreed better with the map. | **Localized reentry** at the posteral septal wall but the ablation target would be at the same location ☑ |
| 23 | CD207 | 648 | 215 | **Macro reentry**: clockwise mitral valve | ✓ |
| 24 | CD208 | 876 | 315 | Hypothesis 1: **Macro reentry** circuit at the septum anterior from the right superior pulmonary vein, active or passive.<br>Hypothesis 2: Same loop as in hypothesis 1, but it might be passive because of another loop with the same pathway that turns around the left veins through the mitral isthmus, coming back over the roof. Ablation at the roof terminated the AT. | ✓ (hypothesis 1) |
| 25 | CD209 | 489 | 270 | **Macro reentry**: roof ascending over the anterior wall and descending over the posterior wall. | ✓ |
| 26 | CD210 | 628 | 340 | **Localized reentry** at the septum | ✓ |
| 27 | CD211 | 603 | 210 | Hypothesis 1: **Localized reentry** or might be larger reentry at the anterior wall near the mitral annulus at 11 o'clock anterior from the LAA.<br>Hypothesis 2: Counterclockwise perimitral circuit: less likely because of colliding wavefront. EP treated as hypothesis 1. | ✓ (hypothesis 1) |
| 28 | CD212a | 1147 | 250 | Hypothesis 1: **Macro reentry** tachycardia roof dependent, ascending on the anterior wall, descending on posterior wall, collision on posterior mitral isthmus.<br>Hypothesis 2: it could be a smaller loop clockwise around perimitral. | ✓ (hypothesis 1) |
| 29 | CD212b | 1089 | 285 | **Macro reentry**: clockwise mitral valve | ✓ |
| 30 | CD213 | 1322 | 305 | There is a SCC at the roof near the RV (there is a gap in a prior ablation line).<br>Hypothesis 1: through SCC a large **macro reentry**: around the RV.<br>Hypothesis 2: through SCC, more localized reentrant circuit using a second gap at the roof for reentry.<br>Hypothesis 3: Both hypothesis might be correct (i.e. double loop).<br>EP ablated at the SCC at the roof, according to hypothesis 1.<br>EP ablated at the SCC at the roof, according to hypothesis 1. | ✓ (hypothesis 1) |
| 31 | CD214 | 439 | 305 | **Macro reentry**: perimitral reentry clockwise pushed away from the mitral annulus (Y shape). | ✓ |

**Table S2.** Percentage of correct cores as a function of increasing noise (i.e. increasing standard deviation *σ* of the Gaussian noise) for varying radius to the true core (0.5 or 1 cm) and a varying number of electrodes (64 or 256). PM stands for phase mapping and DG for DG-mapping. DG-mapping outperforms phase mapping except for a radius of 0.5 cm with 64 electrodes.

| $\sigma$ | 64 | | | | 256 | | | |
| --- | --- | --- | --- | --- | --- | --- | --- | --- |
|  | 0.5 |  | 1 |  | 0.5 |  | 1 |  |
|  | PM | DG | PM | DG | PM | DG | PM | DG |
| 0 | 100 | 100 | 100 | 100 | 100 | 100 | 100 | 100 |
| 5 | 73 | 45 | 92 | 96 | 68 | 98 | 74 | 100 |
| 10 | 46 | 40 | 75 | 91 | 49 | 95 | 62 | 100 |
| 15 | 26 | 35 | 51 | 83 | 19 | 83 | 30 | 95 |
| 20 | 9 | 27 | 21 | 68 | 0 | 62 | 1 | 81 |
| 25 | 2 | 18 | 6 | 50 | 0 | 40 | 0 | 60 |
| 30 | 1 | 12 | 2 | 34 | 0 | 24 | 0 | 41 |

**Fig. S1.**
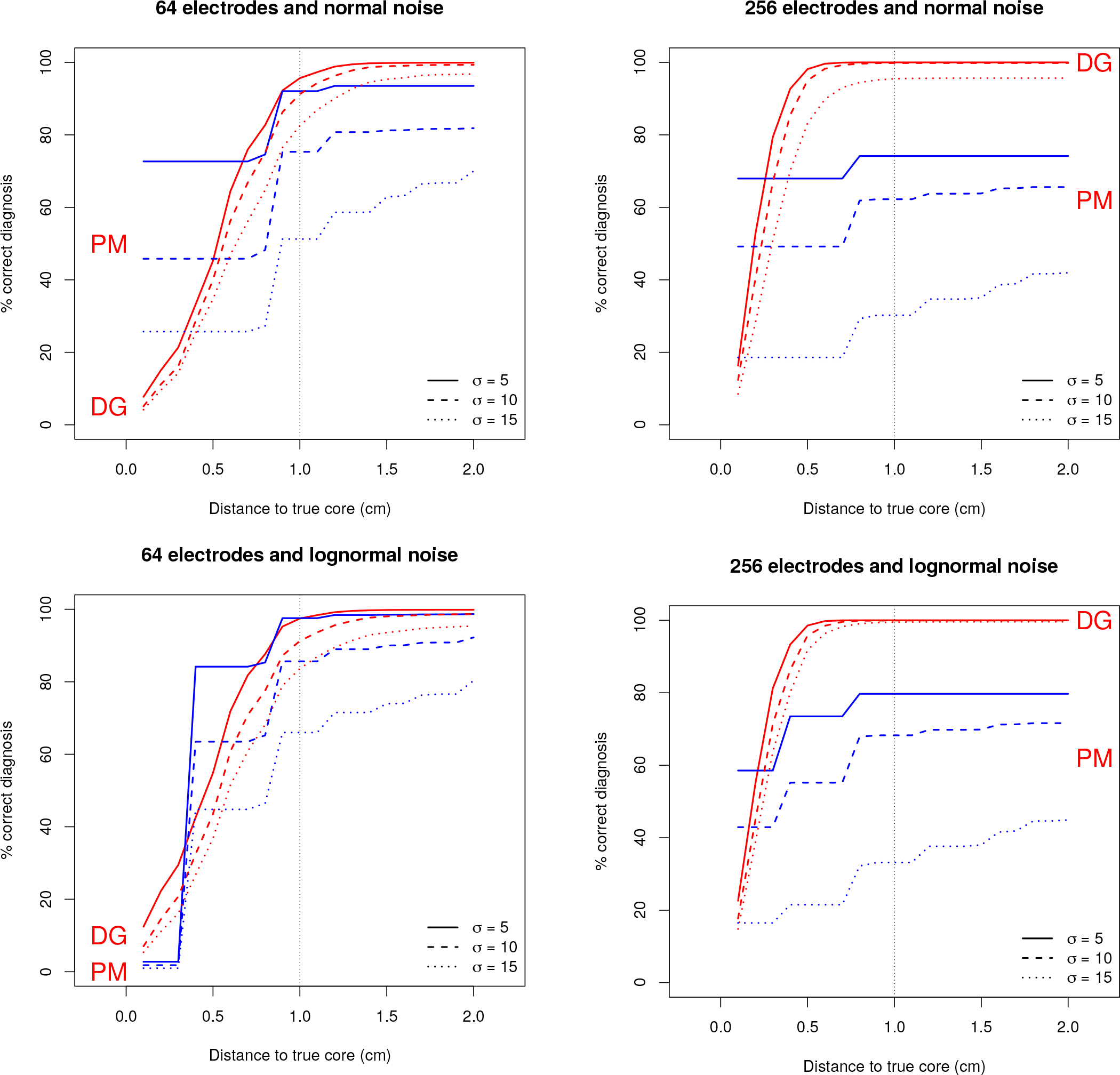
Percentage of correct diagnoses as a function of distance to the true core for different levels of the standard deviation of the noise *σ*. The red lines denote DG-mapping the blue lines phase mapping (PM). The top panels show the performance for normal noise, while for the bottom panels the noise follows a lognormal distribution. DG-mapping has a continuous behavior, while phase mapping shows a stepwise behavior. This is because phase mapping has a resolution determined by the size of the smallest square in the tissue (16×16 mm for the 64 electrode grid, 8×8 mm for the 265 electrode). The minimal steps in performance are seen each 8 mm for 64 electrodes and 4 mm for 265 electrodes, which agrees with the difference of the center of one square and the average of the centers of two contiguous squares, as shown in Figure S2. Therefore, the minimal step in accuracy for phase mapping is determined by the grid size. Phase mapping and DG-mapping can only be compared above this minimal accuracy. The graphs display a superior performance of DG-mapping over phase mapping for both normal and lognormal noise, for all choices of *σ* and for distances to the true core exceeding 1 cm. Overall, the performance of DG-mapping is similar for normal and lognormal noise and improves with increasing number of electrodes.

**Fig. S2.**
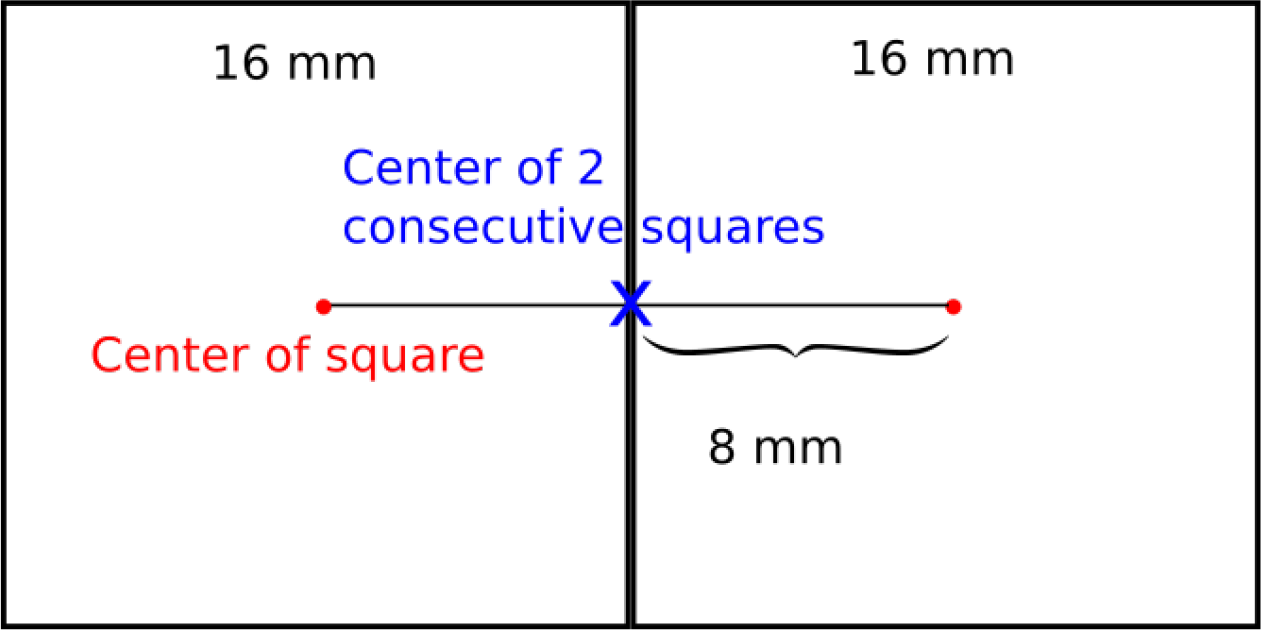
Minimal distance between the center of a square and two consecutive squares for 64 electrodes.

